# Discovering search behaviour in black garden ant trajectories

**DOI:** 10.1101/2025.06.25.661660

**Authors:** Perrine Bonavita, Marius Albino, Jacques Gautrais, Vincent Fourcassié, Maud Combe, Simon Eibner, Christian Jost

## Abstract

Exploration of space plays an important role in many animals, in particular for social insects who have to feed and protect a whole colony. In a laboratory study, Khuong *et al* (2013) studied how the workers of the black garden ant *Lasius niger* move around in an unknown environment. They assumed that, in a homogeneous arena with no visual cues, ants had no information about their position in space. Based on this hypothesis, they modelled their ants in a Boltzmann Walker framework which describes an ant’s random walk as a series of straight segments separated by reorientation events. They assumed that on plain horizontal surfaces the ant’s direction of movement would not influence their average speed, segment lengths and reorientation decisions, thus leading to diffusive trajectories.

However, published experiments indicate that *L. niger* ants are not completely devoid of directional information even in standard laboratory setups with no obvious landmarks. Moreover, many ant species are known to develop specific search strategies when they want to find a particular place in space, a situation that may apply to the analysed data. We re-analyze Khuong *et al* ‘s data on non-inclined surfaces, this time taking into account the ant’s orientation in relation to its starting point in the arena.

We discovered that, with this information taken into account, the ant’s trajectory is biased towards its starting point (biased random walk), revealing an advecto-diffusive process. In fact, the distributions of segment lengths and reorientation angles turned out to be modulated by the ant’s orientation in relation to its starting point. By simulating these biased trajectories, we show that this modulation halves the time it takes for an ant to come back towards its starting point.

We conclude that not taking into account the animal’s cognitive abilities in data analysis may lead to incomplete or biased conclusions. The discovered search behaviour in *L. niger* can play a significant role in the colony’s exploration and foraging ecology.

## Introduction

All organisms move around, either actively or passively, regularly or at specific life stages [1]. Movement, defined as a temporal change of the spatial location of the whole organism, is motivated by diverse processes acting at various spatial and temporal scales [2]. For example, at a large scale animals can perform seasonal migrations linked to changing environmental conditions, while at a smaller scale they can move during foraging, predator avoidance or the search for a partner or a shelter [3].

The study of movement asks several fundamental questions, notably why, how, when and where to move [3]. On longer time scales one may also question the ecological and evolutionary consequences of movement [2]. In an earlier paper, Khuong *et al* (2013) addressed the “how” question while studying the exploratory movement of workers of the individual black garden ant *Lasius niger* in an unknown environment [4]. By studying ant trajectories on flat surfaces of various inclinations (called inclines, including a horizontal incline of zero inclination), their aim was to characterize the influence of gravity on their movement. *L. niger* ants were chosen because they only lay pheromone trails when coming back to their nest from a food source [5]. Therefore, during exploration these ants do not rely on pheromone trails and move independently. The trajectories were modeled within the Boltzmann walker framework (also known as correlated random walks [6–8]) based on a mesoscopic scale [9, 10]. According to this theoretical framework, a trajectory is modeled as a series of straight segments of variable length *λ* (along which the ants move with speed *v*) with turning angles *ω* between them (see Fig 1). The statistical description of the segment length, associated speed and turning angle distributions, permits to characterize the trajectories. In particular, it allows to assess how the trajectories change as a function of inclination and ant orientation *ϕ* (assumed to be the same as segment orientation) with respect to the gradient (direction of steepest slope). This Boltzmann walker model correctly predicted directional dispersal of ants on the inclines [4].

**Fig 1.**
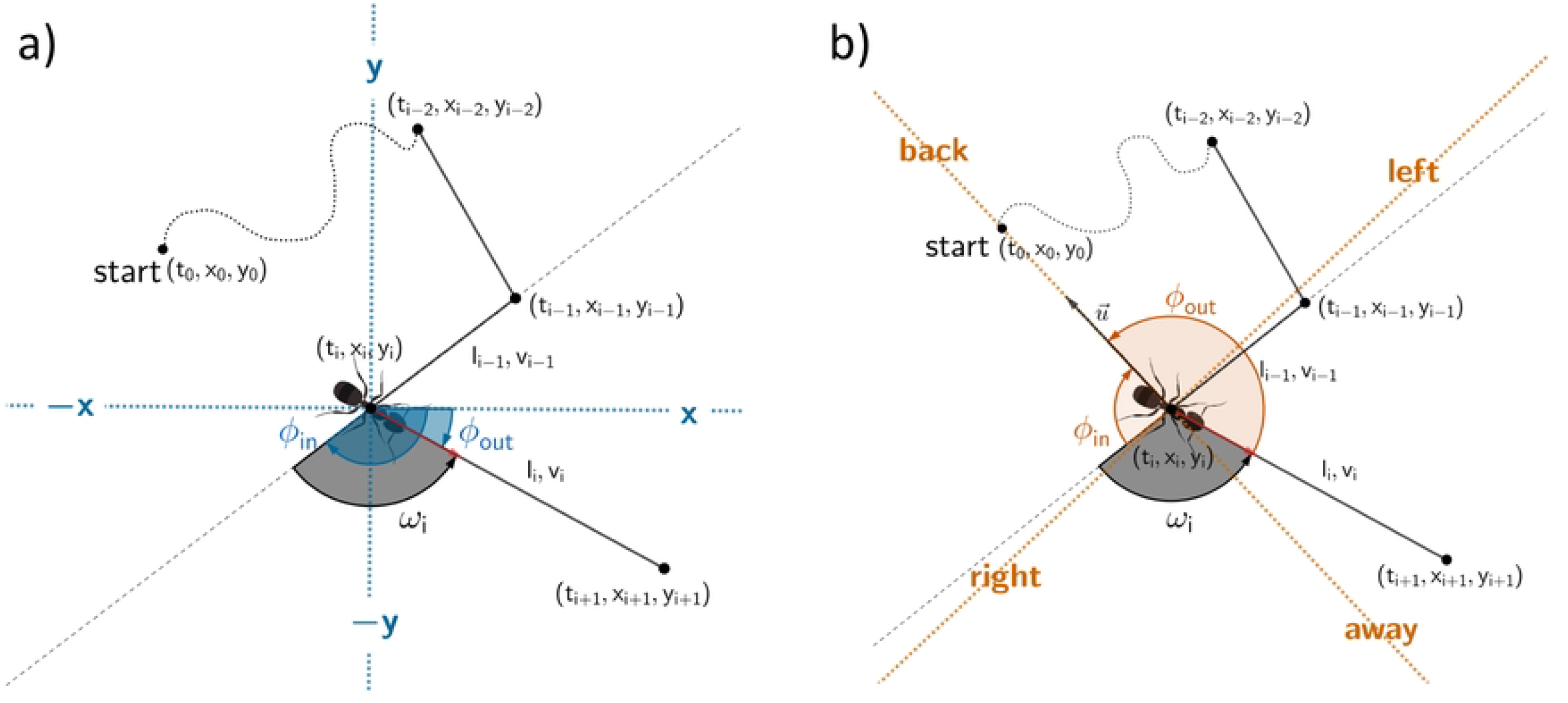
Example of an ant trajectory represented in the Boltzmann walker framework as a series of straight segments of length *li* (along which the ant moves with speed *v*_*i*_) with turning angles *ω*_*i*_ between two successive segments. Ant orientation can be either measured with respect to the x-axis, method used in Khuong’s article (2013) which we will call here the Φ_*x*_ approach (blue (a)), or with respect to the direction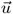 towards the starting point, called 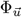 approach (orange (b)). If the ant is currently at position (*t*_*i*_, *x*_*i*_, *y*_*i*_) then *ϕ*_*in*_ refers to the orientation of the incoming segment and *ϕ*_*out*_ to that of the outgoing segment. They thus differ between the two approaches: 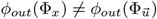 and 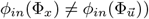. In both approaches, the turning angle is *ω*_*i*_ from *ϕ*_*in*_ to *ϕ*_*out*_. Each trajectory is analyzed with the following model in mind: *v*_*i*_ *∼ v*(*ϕ*_*out*_), *li ∼ l*(*ϕ*_*out*_) and *ω*_*i*_ *∼ ω*(*ϕ*_*in*_).

The trajectory coordinates were obtained in [4] from videos monitoring ant movement on the inclines. With zero inclination the authors assessed ant orientation with respect to the arbitrary *x*-axis of the monitoring camera (what we call the Φ_*x*_ approach in Fig 1(a)). They found, not too surprisingly, that the random walk parameters did not depend on this arbitrary orientation: *v, l*, and *ω* were isotropic with respect to orientation *ϕ*, see Fig 5 in [4] and Fig. 2 in this paper).

**Fig 2.**
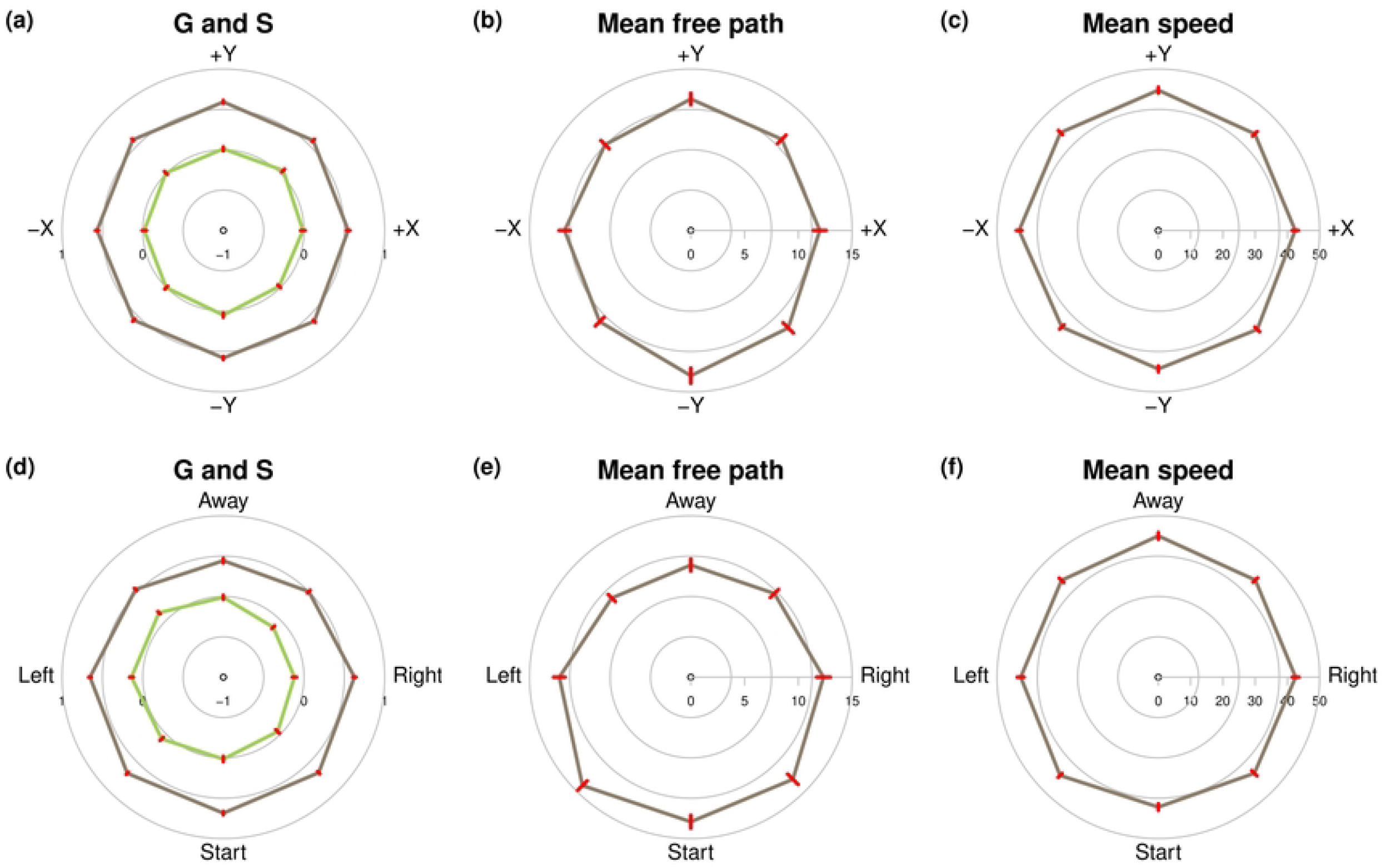
Estimated values of the four Boltzmann random walker parameters *g*_*ω*_, *s*_*ω*_, *λ*, and *v*, as a function of the heading angle with respect to the Φ_*x*_ approach (a,b,c), and the heading angle with respect to 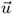 pointing to the starting point, 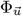 approach, (d,e,f) The distance of a point to the centre of the circles represents the mean parameter value (*±*se) over *n* = 69 trajectories according to the mean orientation of each sector. (a,d) Mean cosine *g*_*ω*_ (red), and mean sine *s*_*ω*_ (green), (b,e) mean segment length *λ* (unit mm), (c,f) mean speed *v* (unit mm/s). Away refers to headings that make the ant move away from the starting point, Start to headings that make it come back towards this point.

When moving in their natural environment, ants are known for their ability to constantly assess their position in space relative to their nest by using a celestial (sun, polarized light) or magnetic compass or memorized visual landmarks [11]. When passively displaced from their current position this cognitive ability allows nestbound ants to perform search behaviour in order to find familiar landmarks to return to their nest [12–16]. It can also be used by ants to search for a renewable food item found at a specific location that ants have memorized in a previous foraging trip [14, 16, 17]. It is characterized by a strong tendency to come back to the point of search initiation and by trajectories forming loops of increasing sizes around this point. The underlying mechanism for search behaviour was found to be the capacity of ants to perform path-integration. Ants are indeed able to integrate constantly the directional and distance components of their trajectory, which means that at all times they know the position of the starting point of their search [18]. Search behaviour has mostly been studied in desert ants of the genus *Cataglyphis* and *Melophorus* but is probably present in most ant species, including in the species *Lasius niger* used in Khuong et al’s (2013) experiment [19].

In general, search behaviour can be formalized as biased correlated random walks [7, 20–22] in which path length, turning angles and speed may depend on a global direction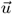 (that may differ between subsequent positions, see Fig. 1(b)). The Boltzmann walker model used by Khuong et al. (2013) [4] can also correspond to such biased random walks for the trajectories on sufaces with non-zero inclinations. Indeed, the authors characterized how the model parameters (form of the turning angle distribution, mean speed, mean segment length) depend on the ant’s orientation with respect to the gradient. In that case, the fixed direction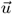 that biased the random walk would correspond to the gradient.

One of the main results of Khuong *et al* ‘s study is that the random walk is not biased on surfaces with zero inclination (the gradient is then zero). However, even with zero inclination, these model variables might depend on the animal’s egocentric position, i.e. its current position and orientation relative to the position of the starting point of its trajectory [23] or any potential landmarks in the environment.

The ants in Khuong et al’s experiments can be considered to be “lost” because they have been passively transported on a wood stick from an exploration point near the nest to an unknown experimental arena. While it is not known whether *L. niger* ants have path-integration capabilities, they can certainly perceive any visual landmarks and laboratory experiments indicate that their speed and U-turn rate can change when they deviate from the path going straight back to the nest, which suggests that they have a knowledge of the spatial location, or at least the direction, of their nest [24].

Do *L. niger* ants also show search behaviour on non-inclined surfaces that can influence trajectories, revealing preferential directions? With this question in mind we re-analyze Khuong et al’s (2013) [4] data on non-inclined surfaces (downloadable from their ESM). However, instead of computing orientation with respect to the arbitrary *x*-axis (subsequently termed the Φ_*x*_ approach, see Figure 1), we now compute orientation with respect to its direction 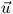 pointing from its current position towards the starting point of the trajectory, which is close to the release location of the ant in the centre of the experimental arena (subsequently termed the 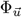 approach, see Figure 1(b)). We reconstruct the distributions of the Boltzmann walker variables *v, l*, and *ω*, by discretizing the orientation angle into 8 sectors (as in [4]) and by assembling the experimental values of the variables in each sector. In contrast to the original analysis in [4] (Φ_*x*_ approach) where they found an isotropic distribution of the distribution parameters of the three variables, we show that with the 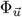 approach privileged directions emerge that support movements typical of search behaviour.

How do *L. niger* ants know their orientation with respect to 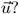 Vision might be involved. To test this hypothesis we ran a series of experiments where ants were monitored either under white light or under red light (which is not perceived by ants). The same trajectory analysis as above should reveal whether search behaviour is reduced or disappears under red-light conditions. Moreover, by using non-parametric trajectory simulations, we assess how reconstructing the distributions according to the 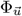 approach, rather than according to the Φ_*x*_ approach, affects the spatio-temporal distribution of ants with respect to their starting point. We will also compute the average time it takes for ants to return towards the starting point, which serves as an indicator of the ecological advantage of search behaviour. Finally, we discuss the link between this modified Boltzmann walker model on non-inclined surfaces (adapted to search behaviour) and macroscopic diffusion type equations.

## Materials and methods

Trajectories on flat non-inclined surfaces were downloaded from the ESM of Khuong *et al*. [4]. The ant species studied and the experimental procedures is described in the paper. There are 69 trajectories. Note that ants were transported on a paint brush from their nest directly to the experimental area (a canvas free of any colony odour), which is the reason why we think that ants will immediately be in a search behaviour mode.

The original trajectories provided in the ESM were sampled at 25 fps. To apply the Boltzmann walker analysis, we adopted the same segmentation algorithm as described in [4]. This segmentation aims at grouping together the positions that belong to the same segment *l*. The algorithm works as follows: initially, the orthogonal distance from each point (*x*_*i*_, *y*_*i*_) to the segment (*x*_*i−*1_, *y*_*i−*1_)–(*x*_*i*+1_, *y*_*i*+1_) is computed for all points. Then, the point with the smallest distance is removed from the trajectory. The distances of the neighboring points of the removed point are then updated, and the process is repeated until the smallest distance found exceeds the threshold *ε* = 1.7 mm. The ant’s timestamps at each point remain unchanged.

The segmented trajectories can now be characterized by the distributions of the three variables used in the Boltzmann walker, *i*.*e. l, v* and *ω*. For each point of the segmented trajectory (except the start and end points), we compute these three quantities, together with the heading angles *ϕ*_*in*_ and *ϕ*_*out*_ for the Φ_*x*_ or the 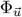 approach (Fig 1). After discretizing the interval [*−π, π*] into 8 angular sectors of the same size with mean values 0, *π/*4, *π/*2,…, we assign each value of *l, v*, and *ω* to the corresponding sector according to the corresponding value of *ϕ* (*ϕ*_*in*_ for *ω, ϕ*_*out*_ for *l* and *v*) for each approach.

Then, for each sector, we computed, over each trajectory, the corresponding estimated value of the mean free path *λ* (estimated as the inverse of the weighted regression slope of the segment length survival curve on a log-linear scale), the median speed *v*_*med*_, and the mean cosine *g*_*ω*_ and mean sine *s*_*ω*_ of the distribution of the turning angles *ω*, defined respectively as the integral of the product of the probability density function Ω for the turning angles *ω* with the cosine and sine functions respectively [22]:

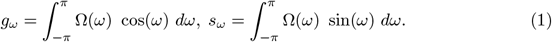

*The valu*es of both *g*_*ω*_ and *s*_*ω*_ statistics lie in the interval [*−*1, 1]. The mean cosine is a measure of the strengths of the persistence in the direction of motion. In symmetric distributions (as expected for the turning angle *ω*), values close to *g*_*ω*_ *≈* 1 correspond to distributions with a peak at *ω ≈* 0, i.e., small turning angles, denoting a high persistence. When turning angles are mostly uniformly distributed, *g*_*ω*_ *≈* 0, and values close to *g*_*ω*_ *≈ −*1 correspond to distributions peaked at large values of *ω*, close to *±π*, as in U-turns. Mean sine *s*_*ω*_ describes the asymmetry of the distribution, with positive values indicating a bias for turning left, and negative values indicating a bias for turning right. Search behaviour may have different impacts on these quantities. One expects for example that when the animal is heading towards the release point, *g*_*ω*_ might be larger than when heading away. Also, when the heading is perpendicular to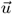 we expect *s*_*ω*_ to be different from 0, biasing turning angles towards 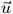.

The standard errors of the mean values, computed over the *N = 69* trajectories, were computed by two methods. For the mean speed that was computed from the median speeds of each trajectory and sector we used the standard formula, i.e., the standard deviation divided by 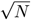 (Fig 2c,f). For the mean free path *λ* and the mean cosine and sine *g*_*ω*_ and *s*_*ω*_ (Fig. 2a,b,d,e), we used a non-parametric bootstrap method [25]. A bootstrap sample was created by first creating 69 bootstrap trajectories by drawing randomly with replacement segment lengths and turning angles from each original trajectory. From these we then drew randomly with replacement a sample of 69 trajectories. From this overall bootstrap sample we computed *λ, g*_*ω*_, and *s*_*ω*_ with the method described above for the original data. This method was repeated 250 times and standard errors were estimated as the standard deviations of these 250 bootstrap parameter estimates.

Assembling all *l, v*, and *ω* values over all original trajectories, either in the Φ_*x*_ or 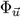 approach, permits us to reconstruct their overall empirical distributions with respect to the orientation sector and inclination. These empirical distributions are used to simulate trajectories non-parametrically to assess the impact of the Φ_*x*_ or 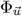 approaches on ant dispersal in a more classical drift-diffusion framework. A simulated trajectory begins at the point (0, 0) with a uniform random orientation, drawing successively segment lengths and turning angles in the empirical distributions corresponding to the orientation sector containing the current ant orientation. In the Φ_*x*_ approach, orientation is computed with respect to the *x*-axis, while in the 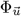 approach we consider a homogeneous vector field where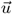 always points in direction of the *−y* axis, as if the ant was far away from its actual starting point. In a first series, trajectories were simulated for 10s to assess the resulting spatial ant distribution. In a second series they were simulated until the ants cross a circle of diameter 200 mm around the origin in order to assess the time it takes them to get that far away. In both cases we simulated a total of 100,000 trajectories to reconstruct either the ant spatial distribution or the time distribution to cross the circle. We compared the times to cross this circle in the two approaches by computing the ratio of the mean time in the Φ_*x*_ approach divided by the mean time in the 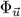 approach. The associated standard error was computed by a non-parametric bootstrap with 100,000 bootstrap samples [25].

To test whether visual cues influence the ant’s search behaviour we run in March 2025 a series of additional experiments similar to those carried out in [4]. We placed ants of three queenless colonies of the same species *L. niger* (captured in October 2024 at Marquefave, France), in a 50 × 45 cm circular arena isolated by black curtains. The experiments took place in a temperature (25°C) and humidity (50%) controlled room. We used the same protocol as in [4]: place a new object in the colony, select with a tooth pick an ant that explores this object, transfer the ant passively on its tooth pick to the arena center and let it descend spontaneously, film its path for 3 minutes or until it reaches the arena wall. We tracked the movement of 60 individual ants, each ant tested under both white light (LED SuperSlim 20W, luminosity 1800-2000 lm or 150 W, from ONSSI) and red light (Lee Filters 787 Marius Red placed in front of the LED lights) conditions (paired design).The light intensity in the arena center was measured using a luxmeter. Under red light the intensity was 17.6 *±* 0.6 lx, while under white light it was 725 *±* 24.5 lx. Colony and light conditions were randomized. After each experiment, the arena was cleaned with alcohol to remove any potential chemical cues [26]. To film the arena from above, we used a Sony ZV-E10 camera with a frame rate of 25 frames per second. These videos were then analysed with a custom-made in-house software, called TOSIA, in order to track the ants and obtain fixed-time trajectories of the individual movements.

The segmentation of the trajectories and their analysis were carried out as described above.

To assess whether individual orientation angles influence linear movement descriptors, we used circular-linear correlation analyses. This method tests for statistical association between a circular variable (here, orientation angles in radians) and a linear variable (e.g., *v, g*_*ω*_, *s*_*ω*_). Correlation strength was quantified using statistic *R*^*2*^, *and s*tatistical significance was assessed using the associated *p-value* [27]. Post-hoc tests were carried out using linear mixed models on simplified segment classes (grouping together the 3 classes at the top and bottom for *v, λ* and *g*_*ω*_, and the 3 classes on the right and left for *s*_*ω*_) in order to be able to identify significant differences while controlling dependencies between segments.

To test the significance of the isotropic or non-isotropic distribution of mean free paths *λ*, we carried out survival analyses on the total data (all the trajectories put together). This methodological choice is justified by the strictly positive and asymmetric nature of the lengths measured, which are similar to survival times in the statistical sense. A log-rank test was used to evaluate the null hypothesis of equality of survival functions between groups, thus revealing whether certain orientations are associated with significantly shorter or longer lengths. A pairwise log-rank test, with Benjamini-Hochberg correction, was used to identify more precisely the pairs of classes showing significant differences.

The trajectory analysis, statistical comparisons and simulations, were all performed with the R software version 4.3 [28] in the RStudio IDE [29]. The corresponding scripts and the segmented trajectories can be requested from the authors. All estimates are reported in the text or the figures as mean *±* standard error.

## Results

Figure 2 summarizes the estimated Boltzmann walker parameters reconstructed as a function of the ant heading angle, either with respect to the *x*-axis (Φ_*x*_ approach) or with respect to the vector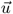 pointing towards the start point (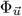 approach). When distributions are built in the Φ_*x*_ approach (Fig. 2a-c), all corresponding parameters look isotropic, there is no effect of orientation on their values (compare to Fig 5 in [4]).

Regarding the mean cosine *g*_*ω*_ and mean sine *s*_*ω*_, the Φ_*x*_ approach (Fig. 2a) yielded a non-significant correlation (respectively *R*^*2*^ *= 0*.*00*31, *p = 0*.*188* and *R*^*2*^ *= 0*.*00*49, *p = 0*.*069*). However, in the 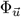 approach (Fig. 2d) we detected a significant correlation (respectively *R*^*2*^ *= 0*.*08*0, *p < 0*.*001* and *R*^*2*^ *= 0*.*01*40, *p < 0*.*001*). An ant coming back to the starting point has a significantly narrower turning angle distribution (*F(1, 200)* = 53, *p < 0*.*001*) (mean cosine *g*_*ω*_ closer to 1: coming back 0.69 *±* 0.02, going away 0.44 *±* 0.03). When it moves perpendicular to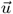, it has a tendency to turn back towards the starting point (mean sine *s*_*ω*_*≠*0, the associated distribution of *ω* is significantly asymmetric (*F(1, 192)* = 61.2, *p < 0*.*001*): left 0.14 *±* 0.03, right *−*0.12 *±* 0.03).

A log-rank test revealed a major significant effect in the 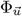 approach (Fig. 2e) of walking direction on the distribution of free path lengths (*χ*^*2*^ *= 279*, df = 7, *p ≪* 0.001). The away direction showed shorter mean free paths (10.4 *±* 0.6 mm) compared to the back direction (13.5 *±* 0.6 mm), indicating a strong anisotropy in movement behaviour (*p ≪* 0.001, pairwise comparisons using Log-Rank test with Benjamini-Hochberg correction). In the Φ_*x*_ approach (Fig. 2b), survival probabilities also differed significantly between the eight orientation classes (log-rank test:

*χ*^*2*^ *= 44*.*4, df = 7, p < 0*.*001)*. This effect on the mean free path length *λ* is mainly due to the slight significant difference between the -Y orientation class and the others: the average free path lengths of ants walking towards -Y are slightly longer (13.4 *±* 0.7 mm) than those of ants walking in other directions, as for example +Y (12.2 *±* 0.7 mm, (*p = 0*.*03*, pairwise comparisons).

For median speed *v*, the Φ_*x*_ approach (Fig. 2c) showed no significant correlation with orientation (*R*^*2*^ *= 0*.*00*02, *p = 0*.*875*), whereas the 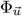 approach (Fig. 2f) revealed a weak but statistically significant correlation (*R*^*2*^ *= 0*.*00*65, *p = 0*.*030*), revealing a slight directional influence on speed in the 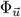 approach. An ant coming back to the starting point moves with a slightly lower speed *v* (40.2 *±* 1 mm/s) than an ant moving away from the starting point (43.8 *±* 1.1 mm/s, (*F(1, 197)* = 8.2, *p = 0*.*005*)).

Fig. 3 shows the results of our own experiments under white light and under red light using the 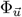 approach.

**Fig 3.**
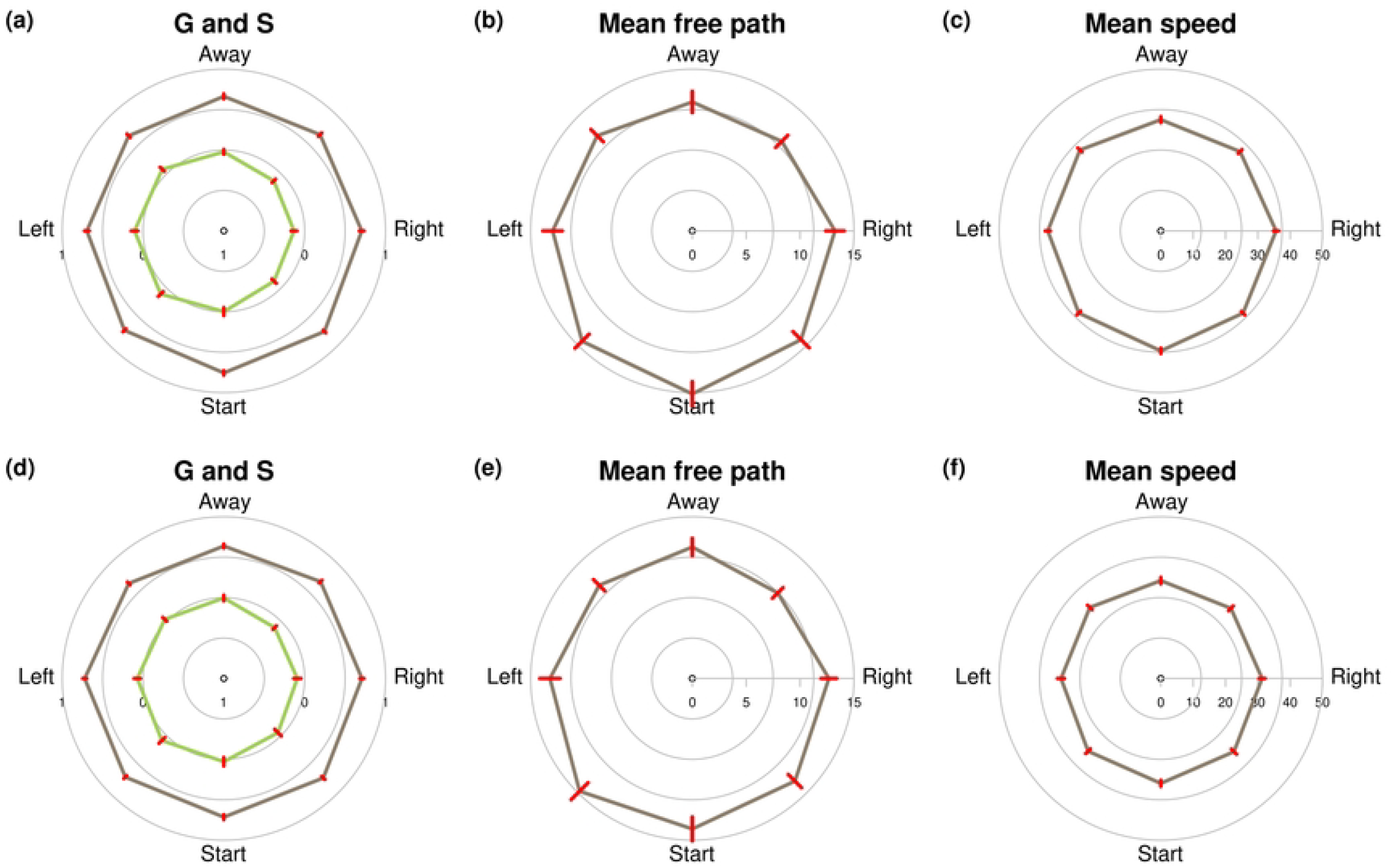
Estimated values of the four Boltzmann random walker parameters *g*_*ω*_, *s*_*ω*_, λ, and *v*, in function of the 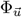 approach (i.e. heading angle with respect to 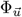 pointing to the starting point), for trajectories under white (a,b,c) and red light (d,e,f) The distance of a point to the centre of the circles represents the mean parameter value (*±*se) over *n* = 60 trajectories according to the mean orientation of each sector. (a,d) Mean cosine *g*_*ω*_ (red), and mean sine *s*_*ω*_ (green), (b,e) mean segment length λ (unit mm), (c,f) mean speed *v* (unit mm/*s*).

Under white light, we found similar results to the data collected in [4]. The ant orientation had a significant effect on the mean free path *λ*, and a log-rank test reveals a significant effect of orientation on the survival curves (*χ*^*2*^ *= 60*.*5*, df = 7, *p ≪* 0.001). The same applies to the median walking speed *v* (*R*^*2*^ *= 0*.*01*0, *p = 0*.*012*). Ants tend to walk longer (15.1 *±* 1.2 mm) and faster (37 *±* 0.8 mm/s) towards their starting point than away from it (11.9 *±* 0.9 mm for *λ*, 34.4 *±* 0.9 mm/s for *v*). The pairwise comparisons using a log-rank test yield *χ = 26*.*7, df = 1, p ≪* 0.001 in the first case and a mixed-Link model on simplified orientation classes yield *F(1, 156)* = 10.7, *p = 0*.*001* in the second. A stronger effect was observed for directional bias: *s*_*ω*_ shows a significant association with orientation (*R*^*2*^ *= 0*.*02*3, *p < 0*.*001*; right: *−*0.13 *±* 0.03 and left: 0.10 *±* 0.04 with *F(1, 156)* = 30, *p ≪* 0.001). In contrast, directional persistence, *g*_*ω*_, was not significantly affected by orientation (*R*^*2*^ *= 0*.*00*4, *p = 0*.*194*).

Under red light, only the median walking speed (*R*^*2*^ *= 0*.*00*83, *p = 0*.*025*6) and mean free path *λ* (*χ*^*2*^ *= 70*.*7*, df = 7, *p ≪* 0.001) were significantly influenced by orientation. Ants tend to walk faster (32.4 *±* 0.9 mm/s) and longer (14.7 *±* 1.1 mm) towards their starting point than away from it (30.2 *±* 1 mm/s with *F(1, 159)* = 6.5, *p = 0*.*01*, 12.1 *±* 0.8 mm with *χ = 13*.*9, df = 1, p ≪* 0.001). No significant effects were detected for the directional persistence *g*_*ω*_ (*R*^*2*^ *= 0*.*00*38, *p = 0*.*187*), or mean sine *s*_*ω*_ (*R*^*2*^ *= 0*.*00*23, *p = 0*.*366*).

To compare the distributions of variables between red and white light conditions, retaining the effect of the orientation factor, we used linear mixed models. It appeared that the ants moved significantly (*χ = 127, df = 1, p << 0*.*001)* faster under white light (35.4 *±* 0.3 mm/s) than under red light (31.4 *±* 0.4 mm/s). With regard to the overall distributions of the parameters *l, g*_*ω*_ and *s*_*ω*_, there was no significant difference depending on whether the individual moved under white or red light. However, with these linear mixed models we find the same evidence of anisotropy for *l* and *s*_*ω*_ that we found earlier with circular comparisons. Therefore, the red condition does not seem to affect the intensity of the anisotropic distribution, it only reduces ant speed.

At this stage, the question arises whether the small observed effects of heading upon the mean cosine *g*_*ω*_, mean sine *s*_*ω*_, mean speed *v*, and mean free path *λ* would be sufficient to yield a change at the level of the trajectories, therefore on ant dispersal. For testing this, and while waiting for a full formalization of the Generalized Boltzmann Walker model embedding the 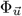 approach, we explore this question numerically with non-parametric path simulations based on the reconstructed empirical parameter distributions (Fig 2). In the Φ_*x*_ approach, ants disperse equally in all directions

(Fig 4a), and the distribution of final positions tends to some gaussian shape, as expected under pure diffusion. By contrast, in the 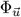 approach, the simulated ants’ final positions show a strong tendency to drift in the direction of 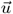 (Fig 4b), as expected under drift-diffusion. Hence, the effect sizes on mean free path, *g*_*ω*_, *s*_*ω*_ and mean speed at the segments’ scale are by far enough to yield a major change at the dispersal scale. This bias also influences the time to move at least 200 mm away from the starting point (Fig 4c), which is 1.92 (*±* 0.02 se) times shorter in the 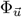 approach compared to the Φ_*x*_ approach.

**Fig 4.**
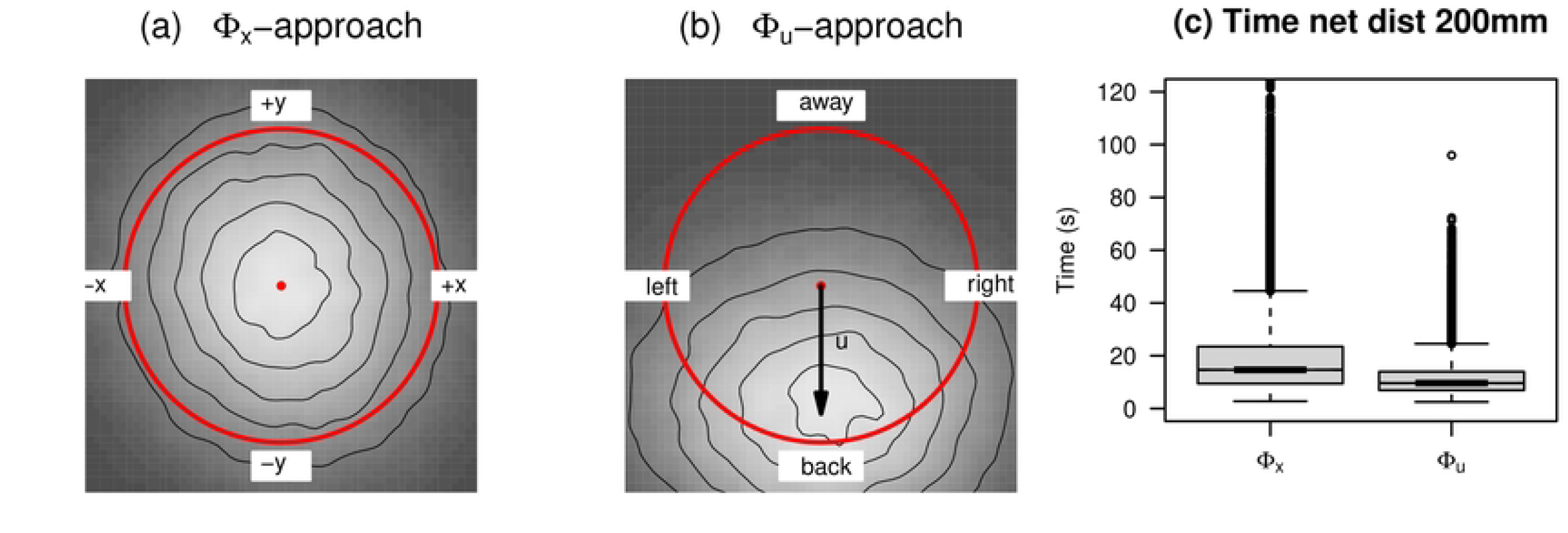
Impact of the Φ_*x*_ (a) and 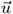 (b) approach on ant movement assessed by non-parametric simulations based on the empirical distributions of *l, v* and *ω* reconstructed either in the Φ_*x*_ or the 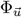 approach. (see M&M for details). (a,b) Distribution (with contour lines) of ants after a 10s path simulated from a point (red central dot) that is far away from the starting point (considered to be in the back direction). White refers to high density, black to low density. In the Φ_*x*_ approach ant orientation is assessed with respect to the *x*-axis, while it is assessed with respect to the 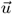 direction in the 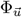 approach. (c) shows the boxplots of times to perform a net displacement of 200 mm for these simulations (red circles in (a,b))

## Discussion

We reanalyzed the movement data on flat homogeneous surfaces from Khuong *et al*. (2013) [4] which led the authors to conclude that the random walk parameters are homogeneous in space, independent of the ant’s orientation. Assuming that these ants know their orientation in space and have a tendency to return to their starting point, we re-estimate these parameters as a function of their orientation with respect to the beginning of their trajectory. We found that these ants perform search behaviour, that is, they have a very strong tendency to remain close to their starting point. The literature already suggests that *L. niger* know where in which direction they move [24], but our analysis is the first firm proof that they can perform search behaviour during their exploration activity, in particular after having been displaced passively to a new area.

The mechanism how they do so is of the taxis type [30]. Their orientation with respect to the starting point influences all parameters involved in a correlated random walk (or Boltzmann walk): ant speed *v* (slightly slower when heading back), free path *λ* (longer when heading back), turning angle parameters mean cosine *g*_*ω*_ (smaller turning angles when heading back) and mean sine *s*_*ω*_ (tendency to turn towards back direction). These findings for Khuong et al’s data are confirmed with our own data set with the same species and in similar conditions, both under white light and under red light. The latter only reduces ant speed and the influence of orientation on mean cosine.

We detected this search behaviour using the same modeling framework as Khuong et al. (the Boltzmann walker or CRW). Other statistical methods exist (see for example the one suggested in [21] and references therein). In future work, we plan to confirm our findings with these other methods. However, as stated in [21], a modeling framework is finally “just a useful means to represent paths in a discrete way”; it has no real meaning for the ant itself. We therefore do not expect our conclusions to change with other methods, though the associated statistical power may be modified.

*L. niger* ants are known to use visual cues to navigate in their environment [31–33]. In our experiment, the area on which ants were moving during the tests was isolated visually from the experimental room with white sheets. However, these sheets did not diffuse light in an homogeneous manner; ants may thus be able to locate the approximate location of the starting point of their trajectory through a matching process between the current view of the arena and a view stored at this point [34]. This may explain how they are able to perform the biased random walk towards this point we observed in our experiment. However, we detected this search behaviour even under red-light conditions, indicating that the ant’s orientation mechanism does not rely solely on vision (or that the red light we used si not completely imperceptible for *L. niger* [35]).

It is not an easy task to obtain general predictions from random walk model formulations (see review in [22]). One solution is the passage to macroscopic equations that are more accessible to mathematical analysis. We have interpreted the Boltzmann walker modeling framework used in Khuong et al (2013) as a correlated random walk (CRW, the found model parameters are isotropic, they do not depend on ant orientation in space). Macroscopic diffusion type models can be derived from this CRW by mesoscopic techniques (see [36], or the appendix in [37]) which also permit to clearly identify any approximations that are made in the process. The case of the CRW suggested by Khuong et al’s analysis leads to the classical telegraph equation if we only make the so-called P1-approximation (applies when the spatial scale of interest requires many turning events [38]), or to the standard Fickian diffusion equation if we also make the diffusion approximation (flux density dynamics can be considered stationary compared to animal density dynamics, which applies when mean free paths are much smaller than the spatial scales of any environmental heterogeneity that influences mean free paths). In both cases the macroscopic model predicts that average ant density evolves as a Gaussian distribution around the starting point, with a standard deviation that is proportional to the square root of time. Ants therefore can move arbitrarily far away from the starting point as time goes on (net squared displacement increases linearly with time [39]). With our detected dependence of model parameters (form of the turning angle distribution, mean speed, mean segment length) on the animal’s orientation with respect to the direction from its current position to its initial location, the Boltzmann walker becomes a biased correlated random walk BCRW (sensu [22], especially their §3). We expect this effect to “constrain” ants close to the starting point (net squared displacement should become sub-diffusive, maybe even bounded). The corresponding macroscopic models (and the approximations made when deriving them from a mesoscopic model) have to be reassessed, in particular their pertinence for a given biological context. The literature mostly address the more general case of drift, that is a bias to move towards a given fixed direction [22, 40]. The macroscopic model becomes in this case a drift-diffusion (or advection-diffusion) equation. The BCRW case of searching behaviour where the bias depends on the animal’s orientation with respect to a fixed position in space seems to be less explored in the literature. A further comprehensive literature search combined with the appropriate mathematical developments will be necessary in order to assess whether such macroscopic models are suitable for ant searching behaviour. In any case, our non-parametric simulations illustrate that subtle changes in random walk parameter values can have a profound impact on ecologically relevant statistics such as return times towards a starting point. Further theoretical investigations, either on the individual based model level or on the macroscopic level (and the passage to get there) will prove useful to better assess such effects for a given experimental or field situation.

In conclusion, our analyses show that *L niger* ants show search behaviour when they find themselves in an unknown environment. They retain this search behaviour even under red-light conditions, showing that vision is not the only sense used to perform search behaviour. At the macroscopic level this indicates that ant dispersal does not proceed as predicted by simple diffusion models, but that advective type dispersal emerges, permitting ants to stay close to their initial position even on long time scales. The impact of this search behaviour on colony level functions will require further investigations.

## Acknowledgments

We would like to thank the researchers and non-permanent members of the CAB team at CRCA and the MesoStar team at LAPLACE for their advice on writing this article. We would also like to thank Loïc Lacour for his precious help to perform and digitize the experiments under white and red light.

## References

1. Nathan R, Monk CT, Arlinghaus R, Adam T, Alós J, Assaf M, et al. Big-data approaches lead to an increased understanding of the ecology of animal movement. Science. 2022;doi:10.1126/science.abg1780.

2. Nathan R, Getz WM, Revilla E, Holyoak M, Kadmon R, Saltz D, et al. A movement ecology paradigm for unifying organismal movement research. Proceedings of the National Academy of Sciences of the United States of America. 2008;doi:10.1073/pnas.0800375105.

3. Swingland IR, Greenwood PJ. The Ecology of Animal Movement. The Journal of Animal Ecology. 1985;doi:10.2307/4647.

4. Khuong A, Lecheval V, Fournier R, Blanco S, Weitz S, Bezian JJ, et al. How Do Ants Make Sense of Gravity? A Boltzmann Walker Analysis of Lasius niger Trajectories on Various Inclines. PLOS ONE. 2013;8(10):e76531. doi:10.1371/journal.pone.0076531.

5. Beckers R, Goss S, Deneubourg JL, Pasteels JM. Trail laying behaviour during food recruitment in the ant Lasius niger (L.). Insectes Sociaux. 1992;39(1):59–72. doi:10.1007/BF01240531.

6. Kareiva PM, Shigesada N. Analyzing insect movement as a correlated random walk. Oecologia. 1983;56(2-3):234–238. doi:10.1007/BF00293798.

7. Marsh LM, Jones RE. The form and consequences of random walk movement models. Journal of Theoretical Biology. 1988;133(1):113–131. doi:10.1016/S0022-5193(88)80028-6.

8. Challet M, Fourcassié V, Blanco S, Fournier R, Theraulaz G, Jost C. A new test of random walks in heterogeneous environments. Naturwissenschaften. 2005;92(8):367–370. doi:10.1007/s00114-005-0001-1.

9. Gautrais J. Habilitation à Diriger des Recherches [PhD Thesis]. Université Toulouse 3 Paul Sabatier; 2015.

10. Khuong A. Modéle comportemental de la dynamique de construction de la structure épigée du nid chez la fourmi Lasius niger: approches expérimentales et théoriques [PhD Thesis]. Université de Toulouse, Université Toulouse III-Paul Sabatier; 2013.

11. Wehner R. Desert navigator: the journey of an ant. Belknap press; 2020.

12. Wehner R, Srinivasan MV. Searching Behaviour of Desert Ants, Genus Cataglyphis (Formicidae, Hymenoptera). Journal of Comparative Physiology A. 1981;doi:10.1007/BF00605445.

13. Müller M, Wehner R. The Hidden Spiral: Systematic Search and Path Integration in Desert Ants, Cataglyphis fortis. Journal of Comparative Physiology A. 1994;175:525–530. doi:10.1007/BF00199474.

14. Fourcassié V, Traniello JFA. Food Searching Behaviour in the Ant Formica schaufussi (Hymenoptera, Formicidae): Response of Naive Foragers to Protein and Carbohydrate Food. Animal Behaviour. 1994;48(1):69–79. doi:10.1006/anbe.1994.1212.

15. Schultheiss P, Cheng K. Finding the Nest: Inbound Searching Behaviour in the Australian Desert Ant, Melophorus bagoti. Animal Behaviour. 2011;doi:10.1016/j.anbehav.2011.02.008.

16. Schultheiss P, Wystrach A, Legge ELG, Cheng K. Information content of visual scenes influences systematic search of desert ants. Journal of Experimental Biology. 2013;216(4):742–749. doi:10.1242/jeb.075077.

17. Pfeffer ES, Bolek S, Wolf H, Wittlinger M. Nest and food search behaviour in desert ants, Cataglyphis: a critical comparison. Animal Cognition. 2015;18(4):885–894. doi:10.1007/s10071-015-0858-0.

18. Heinze S, Narendra A, Cheung A. Principles of Insect Path Integration. Current Biology. 2018;28(17):R1043–R1058. doi:10.1016/j.cub.2018.04.058.

19. Le Breton J, Fourcassié V. Information transfer during recruitment in the ant Lasius niger L. (Hymenoptera: Formicidae). Behavioral Ecology and Sociobiology. 2004;55(3):242–250. doi:10.1007/s00265-003-0704-2.

20. Harkness RD, Maroudas NG. Central Place Foraging by an Ant (Cataglyphis bicolor Fab.): A Model of Searching. Animal Behaviour. 1985;33(3):916–928. doi:10.1016/S0003-3472(85)80026-9.

21. Benhamou S. Detecting an Orientation Component in Animal Paths When the Preferred Direction is Individual Dependent. Ecology. 2006;87:518–528. doi:10.1890/05-0495.

22. Codling EA, Plank MJ, Benhamou S. Random Walk Models in Biology. Journal of The Royal Society Interface. 2008;5(25):813–834. doi:10.1098/rsif.2008.0014.

23. Benhamou S, Sauvé JP, Bovet P. Spatial Memory in Large Scale Movements: Efficiency and Limitation of the Egocentric Coding Process. Journal of Theoretical Biology. 1990;145(1):1–12. doi:10.1016/S0022-5193(05)80531-4.

24. Beckers R, Deneubourg JL, Goss S. Trails and U-turns in the Selection of a Path by the Ant Lasius niger. Journal of Theoretical Biology. 1992;159(4):397–415. doi:10.1016/S0022-5193(05)80686-1.

25. Efron B, Tibshirani RJ. An Introduction to the Bootstrap. 1st ed. Chapman and Hall/CRC; 1994. Available from: 10.1201/9780429246593.

26. Devigne C, Detrain C. Collective exploration and area marking in the ant Lasius niger. Insectes Sociaux. 2002;49:357–362.

27. Fisher NI. Statistical Analysis of Circular Data. Cambridge New York: Cambridge University Press; 1993.

28. R Core Team. R: A Language and Environment for Statistical Computing; 2025. Available from: https://www.R-project.org/.

29. Posit team. RStudio: Integrated Development Environment for R; 2025. Available from: http://www.posit.co/.

30. Fraenkel GS, Gunn DL. The Orientation of Animals: Kineses, Taxes and Compass Reactions. New York: Dover Publications, Inc.; 1961.

31. Aron S, Beckers R, Deneubourg JL, et al. Memory and Chemical Communication in the Orientation of Two Mass-Recruiting Ant Species. Insectes Sociaux. 1993;40:369–380. doi:10.1007/BF01253900.

32. Evison SEF, Petchey OL, Beckerman AP, et al. Combined Use of Pheromone Trails and Visual Landmarks by the Common Garden Ant Lasius niger. Behavioral Ecology and Sociobiology. 2008;63:261–267. doi:10.1007/s00265-008-0657-6.

33. Grüter C, Maitre D, Blakey A, Cole R, Ratnieks FLW. Collective Decision Making in a Heterogeneous Environment: Lasius niger Colonies Preferentially Forage at Easy to Learn Locations. Animal Behaviour. 2015;104:189–195. doi:10.1016/j.anbehav.2015.03.017.

34. Wystrach A, Beugnon G, Cheng K. Landmarks or Panoramas: What Do Navigating Ants Attend to for Guidance? Frontiers in Zoology. 2011;8:21. doi:10.1186/1742-9994-8-21.

35. Yilmaz A, Spaethe J. Colour vision in ants (Formicidae, Hymenoptera). Philosophical Transactions of the Royal Society B. 2022;377:20210291. doi:10.1098/rstb.2021.0291.

36. Patlak CS. Random Walk with Persistence and External Bias. Bulletin of Mathematical Biophysics. 1953;15:311–338. doi:10.1007/BF02476407.

37. Casellas E, Gautrais J, Fournier R, Blanco S, Combe M, Fourcassié V, et al. From Individual to Collective Displacements in Heterogeneous Environments. Journal of Theoretical Biology. 2008;250:424–434.

38. Case K, Zweifel P. Linear Transport Theory. New York, USA: Addison-Wesley; 1967.

39. Einstein A. über die von der molekularkinetischen Theorie der Wärme geforderte Bewegung von in ruhenden Flüssigkeiten suspendierten Teilchen. Annalen der Physik. 1905;17:549–560.

40. Codling EA, Bearon RN, Thorn GJ. Diffusion about the mean drift location in a biased random walk. Ecology. 2010;91(10):3106–3113. doi:10.1890/09-1729.1.

